# Genomic characterization of metastatic patterns from prospective clinical sequencing of 25,000 patients

**DOI:** 10.1101/2021.06.28.450217

**Authors:** Bastien Nguyen, Christopher Fong, Anisha Luthra, Shaleigh A. Smith, Renzo G. DiNatale, Subhiksha Nandakumar, Henry Walch, Walid K. Chatila, Ramyasree Madupuri, Ritika Kundra, Craig M. Bielski, Brooke Mastrogiacomo, Adrienne Boire, Sarat Chandarlapaty, Karuna Ganesh, James J. Harding, Christine A. lacobuzio-Donahue, Pedram Razavi, Ed Reznik, Charles M. Rudin, Dmitriy Zamarin, Wassim Abida, Ghassan K. Abou-Alfa, Carol Aghajanian, Andrea Cercek, Ping Chi, Darren Feldman, Alan L. Ho, Gopakumar Iyer, Yelena Y. Janjigian, Michael Morris, Robert J. Motzer, Eileen M. O’Reilly, Michael A. Postow, Nitya P. Raj, Gregory J. Riely, Mark E. Robson, Jonathan E. Rosenberg, Anton Safonov, Alexander N. Shoushtari, William Tap, Min Yuen Teo, Anna M. Varghese, Martin Voss, Rona Yaeger, Marjorie G. Zauderer, Nadeem Abu-Rustum, Julio Garcia-Aguilar, Bernard Bochner, Abraham Hakimi, William R. Jarnagin, David R. Jones, Daniela Molena, Luc Morris, Eric Rios-Doria, Paul Russo, Samuel Singer, Vivian E. Strong, Debyani Chakravarty, Lora H. Ellenson, Anuradha Gopalan, Jorge S. Reis-Filho, Britta Weigelt, Marc Ladanyi, Mithat Gonen, Sohrab P. Shah, Joan Massague, Jianjiong Gao, Ahmet Zehir, Michael F. Berger, David B. Solit, Samuel F. Bakhoum, Francisco Sanchez-Vega, Nikolaus Schultz

## Abstract

Progression to metastatic disease remains the main cause of cancer death. Yet, the underlying genomic mechanisms driving metastasis remain largely unknown. Here, we present MSK-MET, an integrated pan-cancer cohort of tumor genomic and clinical outcome data from more than 25,000 patients. We analyzed this dataset to identify associations between tumor genomic alterations and patterns of metastatic dissemination across 50 tumor types. We found that chromosomal instability is strongly correlated with metastatic burden in some tumor types, including prostate adenocarcinoma, lung adenocarcinoma and HR-positive breast ductal carcinoma, but not in others, such as colorectal adenocarcinoma, pancreatic adenocarcinoma and high-grade serous ovarian cancer. We also identified specific somatic alterations associated with increased metastatic burden and specific routes of metastatic spread. Our data offer a unique resource for the investigation of the biological basis for metastatic spread and highlight the crucial role of chromosomal instability in cancer progression.

## INTRODUCTION

Although most cancer deaths are due to metastatic spread, little is known about the genomic determinants of cancer metastasis. Once metastatic cancer cells have detached from the primary tumor site, they can invade all parts of the body (Lambert et al., 2017; Massagué and Obenauf, 2016). However, the distribution of metastatic sites for a given primary tumor is not random and is dictated by factors such as anatomical location, cell of origin and molecular subtype, among others (Gao et al., 2019; Nguyen et al., 2009). Furthermore, tumor cell-extrinsic factors such as treatment, target organ microenvironment and other systemic factors such as circulating chemokines and cytokines can also influence the pattern of metastatic progression (Massagué and Ganesh, 2021). The classical seed-and-soil hypothesis, according to which disseminated cancer cells preferentially colonize organs that enable and are compatible with their own growth, has been explored for more than a century (Paget, 1889). Yet much remains unknown about the interplay between tumor genomic features and metastatic potential, as well as organ-specific patterns of metastasis.

Molecular profiling of tumors coupled with clinical annotation of metastatic events could help provide insight into this question. However, large-scale cancer sequencing efforts have so far focused on primary, untreated tumors (e.g., The Cancer Genome Atlas (Sanchez-Vega et al., 2018)), or they have characterized the overall genomic landscape of metastatic disease without explicitly interrogating specific routes of metastatic dissemination (Priestley et al., 2019; Robinson et al., 2017; Zehir et al., 2017). Other studies have investigated the genomic complexity of cancer metastasis by reconstructing tumor evolution across different organs at varying levels of resolution, but they have been limited by small sample sizes (Brastianos et al., 2015; Brown et al., 2017; Eckert et al., 2016; Hu et al., 2020; Jimenez-Sanchez et al., 2017; Makohon-Moore et al., 2017; Naxerova et al., 2017; Noorani et al., 2020; Reiter et al., 2020; Shih et al., 2020). Identifying associations between genomic features and specific patterns of metastatic spread is an active area of research and several landmark studies on this topic have been published during the past few years (Birkbak and McGranahan, 2020). In particular, richly annotated datasets combining genomic features and detailed clinical history of metastases for individual patients have been recently made available through large collaborative efforts such as METABRIC in breast cancer (Rueda et al., 2019) and TRACERx in clear-cell renal cell carcinoma (Turajlic et al., 2018). However, a study involving thousands of participants across multiple tumor types in which clinical and genomic data has been homogeneously processed through a unified computational pipeline is still lacking.

We assembled a pan-cancer cohort of >25,000 patients with tumor genomic profiling and clinical information on metastatic events and outcomes, which we designate MSK-MET (Memorial Sloan Kettering - Metastatic Events and Tropisms). All samples were profiled using the MSK-IMPACT targeted sequencing platform (Cheng et al., 2015), which identifies somatic mutations, rearrangements and copy-number alterations in 341-468 cancer genes, as well as tumor mutational burden (TMB), chromosomal instability and microsatellite instability. Metastatic events were extracted from the electronic health records (EHR) and mapped to a reference set of 21 anatomic locations. We analyzed genomic differences between primary and metastatic samples and between primary tumors from metastatic and non-metastatic patients, stratified by tumor type and molecular subtypes. Our analysis identified associations between metastatic burden (defined as the number of distinct organs affected by metastases throughout a patient’s clinical course) and specific genomic features, including mutational burden, chromosomal instability, and somatic alterations in individual cancer genes. We also identified associations between genomic alterations and organ-specific patterns of metastatic dissemination and progression. The clinical and genomic data used in our study have been made publicly available and constitute a valuable resource that will help further our understanding of metastatic disease.

## RESULTS

### Overview of the MSK-MET cohort

A total of 25,775 patients were included in the present study, consisting of 15,632 (61%) primary and 10,143 (39%) metastatic specimens spanning 50 different tumor types (**Figure S1A-D; Table S1**). The median interval between sample acquisition and sequencing was 62 days (interquartile range (IQR) = 0-287 days). The median sequencing coverage was 653x (IQR = 525-790x) and the median tumor purity assessed by pathologists was 40% (IQR = 20-50%) (**Figure S1B**). The majority of sequenced samples obtained from metastatic sites were from lymph nodes (n=2305, 23%), liver (n=2289, 23%), lung (n=982, 10%), or bone (n=726, 7%). Among primary tumors, 11,741 (75%) were from patients with metastatic disease at the time of sequencing or at a later time (**Figure S1D**). Over the entire course of the disease, a total of 99,419 metastatic events from 21,546 metastatic patients were retrieved from the EHR and mapped to 21 organ sites. The most common target organ sites were lung, liver, bone, or unspecified (**Figure S1E**). The frequencies of organ-specific metastasis of individual tumor types were similar to previous reports (Budczies et al., 2015; Gao et al., 2019) (**Figure S1F**). Internal validation using 4,859 (22.5%) patients included in previous studies with available metastatic events extracted through manual chart review (Abida et al., 2017; Jones et al., 2021; Razavi et al., 2018; Shoushtari et al., 2021; Yaeger et al., 2018) revealed a high concordance and sensitivity with metastatic events extracted from the EHR (**Figure S1G-H**). We used this data to map patterns of metastatic dissemination from 50 tumor types to 21 metastatic organ sites (**Figure 1**).

**Figure 1.**
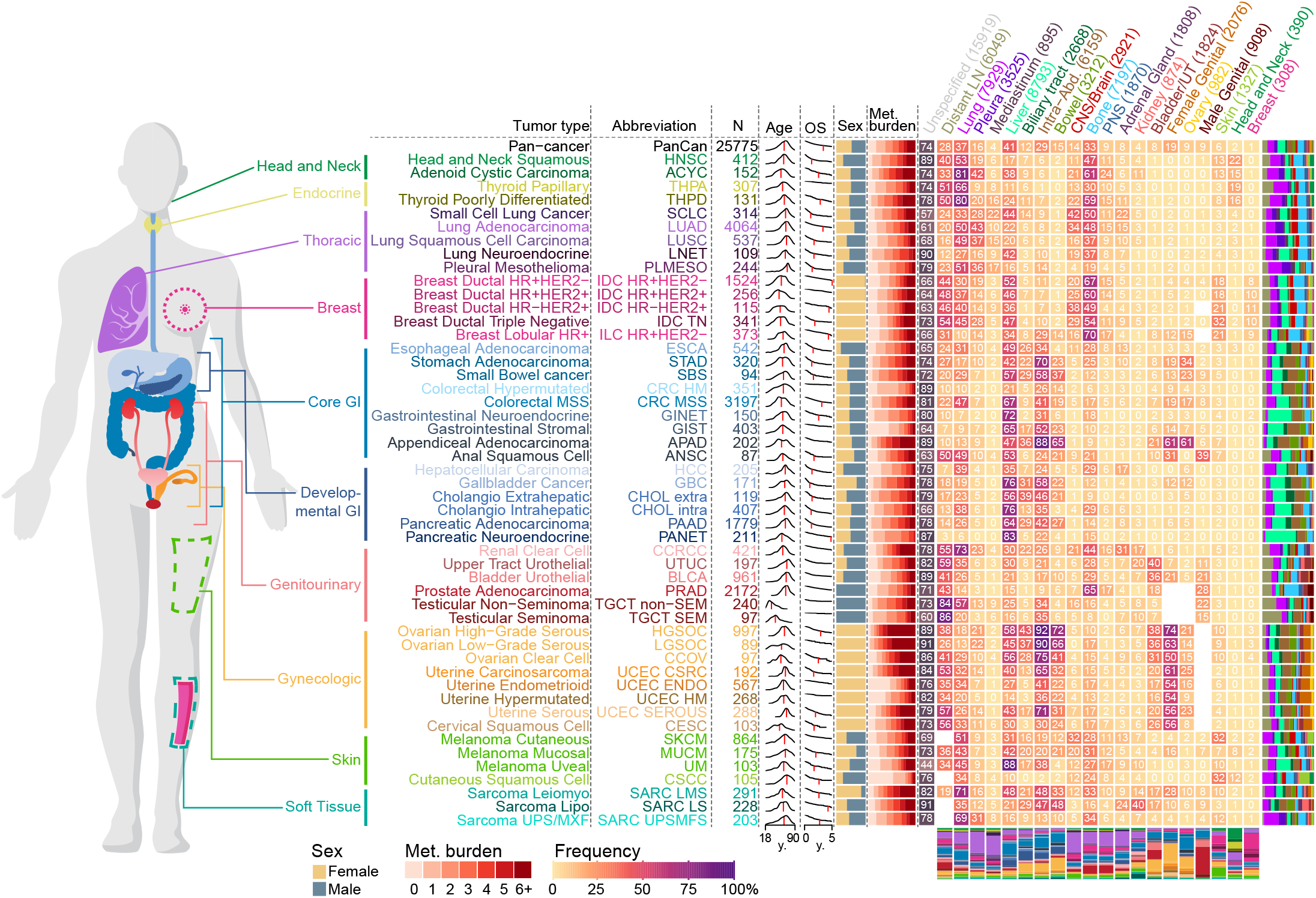
Overview of the MSK-MET cohort. Metastatic patterns of 50 tumor types. For each tumor type, the following attributes are shown from left to right: tumor type abbreviation, number of patients, distribution of age at sequencing (red vertical line indicates the median), overall survival in years from time of sequencing (red vertical line indicates the median OS), sex ratio (female = gold, male = grey), distribution of metastatic burden across all patients (ranging from 0 to ≥6 distinct metastatic sites), and a heatmap with the percentage of metastatic patients with metastases at specific metastatic sites (the entire clinical course was taken into consideration). The number in each cell indicates the frequency of patients having at least one reported metastasis at that given site. For each tumor type, the distribution of all metastasis events by 21 organ sites is shown as a stacked barplot to the right of the heatmap. For each metastatic site, the distribution of all 50 tumor types is shown as a stacked barplot below the heatmap. For each metastatic site, the number of patients having at least one metastasis is indicated in parentheses. Frequencies for sex-specific target organs (female genital, ovary and male genital) were calculated using patients of the corresponding sex. See also Table S1 and Figure S1.

For the whole cohort, the median age at sequencing was 64y, ranging from a median of 33y for patients with testicular non-seminoma to a median of 70y for patients with cutaneous squamous cell carcinoma. Overall, the median follow-up time was 30 months and the five-year survival rate was 40%, ranging from 90% in testicular seminoma to 10% in pancreatic adenocarcinoma. There was a median of four metastatic events per patient, ranging from one in hypermutated colorectal cancer to eight in high-grade serous ovarian carcinoma. Metastatic patterns differed by tumor types and histological subtypes. For example, compared to lung adenocarcinoma, lung neuroendocrine cancer had a higher prevalence of liver metastasis but a lower prevalence of CNS/Brain metastasis. Similarly, lobular breast cancer had a lower prevalence of lung metastasis but a higher prevalence of bone, ovary and peritoneum metastasis, compared to ductal breast cancer as reported before (Borst and Ingold, 1993). Differences in metastatic patterns were also observed across molecular subtypes of the same tumor type. For example, and in line with a previous study (Kennecke et al., 2010), HR-/HER2+ ductal breast cancer had a higher prevalence of CNS/Brain metastasis, while the HR+/HER2- subtype had a higher prevalence of bone metastasis.

### Genomic differences between primary and metastatic tumors

To determine sample type specific genomic differences across 50 tumor types, we compared the genomic features of primary (n=15,632) and metastatic tumors (n=10,143) (independent of the metastatic status of patients). The number of sequenced primaries was higher than the number of sequenced metastases for most tumor types, with some exceptions, such as cutaneous melanoma, high-grade serous ovarian cancer and adenoid cystic carcinoma. In 16 tumor types, metastases were significantly more chromosomally unstable, as inferred by a higher fraction of genome altered (FGA), compared to primary tumors, consistent with previous findings (Bakhoum et al., 2018; Ben-David and Amon, 2020; Hieronymus et al., 2018; Shukla et al., 2020; Stopsack et al., 2019; Watkins et al., 2020) (**Figure 2A-B; Table S2**). The difference in tumor purity- and ploidy-adjusted FGA (adjusted FGA) was confirmed in 11 tumor types using a subset of samples with available FACETS data (n=17,224) (**Table S2**). FACETS allowed us to estimate the frequency of whole-genome duplication (WGD) and assess the clonality of individual variants. As previously reported, WGD frequencies varied across tumor types (Bielski et al., 2018). In eight tumor types, we observed a significantly higher frequency of WGD in metastases compared to primary tumors (**Figure 2A-B; Table S2**). The higher chromosomal instability and higher frequency of WGD were particularly marked in uterine endometrioid, which can be explained by differences in the distribution of genomic subtypes within these two groups (Cancer Genome Atlas Research Network et al., 2013). Tumor mutational burden (TMB) was significantly higher in metastases from 10 tumor types, while TMB was lower only in metastases from hypermutated uterine cancer (**Figure 2A-B; Table S2**). Consistent with the evolutionary bottleneck hypothesis (Birkbak and McGranahan, 2020), metastases from 12 tumor types were significantly more homogeneous, with a higher fraction of clonal mutations compared to primary tumors.

**Figure 2.**
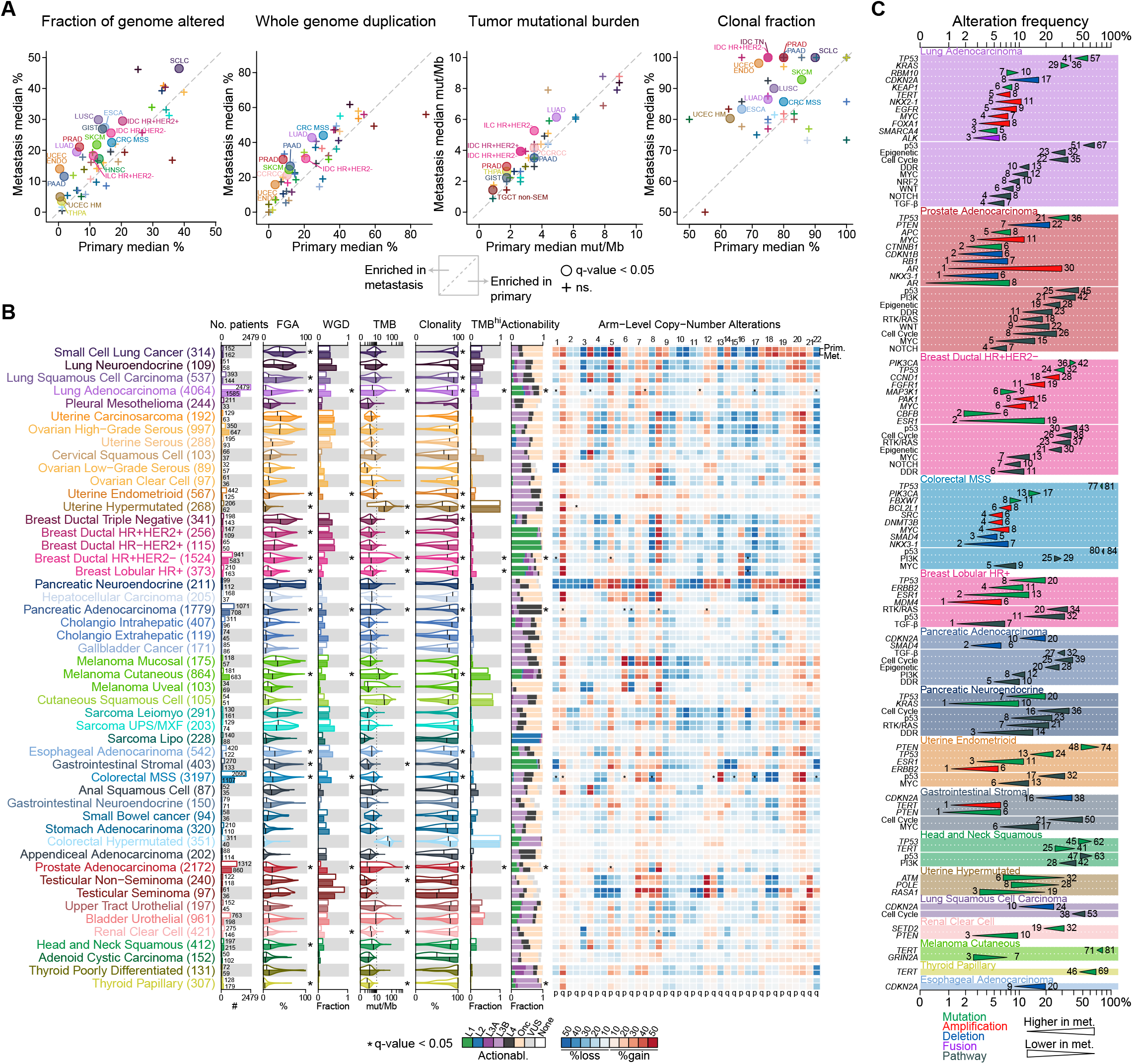
Genomic differences between primary tumors and metastases. (A) Comparisons of the median fraction genome altered (FGA), median whole-genome duplication (WGD) frequency, median tumor mutation burden (TMB), and median clonal fraction for each tumor type in metastatic vs. primary tumors. Tumor types with statistically significant differences are labeled. For TMB both axes were limited to 10mut/Mb. (B) The following clinical and genomic features are shown side-by-side for primary (top row within each cancer type) and metastatic (bottom row) sequenced samples using a combination of barplots and violin plots; from left to right: sample counts, FGA, fraction of samples with WGD, TMB, clonality, fraction of samples with high TMB, and distribution of the highest actionable alteration levels. The black vertical line in each violin plots represents the median. The heatmap shows the frequency of individual arm level alterations in primary tumors and metastases (only the frequency of the more frequent event, gain or loss, is shown). Tumor types are ordered from top to bottom by decreasing FGA in metastasis and grouped by organ systems. * indicates q-value < 0.05. WGD and clonality were available for a subset of 17,224 samples with FACETS data. (C) Statistically significant differences in the frequency of oncogenic alterations and pathways between primary tumors and metastases in individual tumor types. Triangles summarize oncogenic alteration frequencies in primary tumors vs. metastases and are colored according to alteration type. Gene names in italics refer to specific genes, those in regular font refer to pathways. See also Table S2.

We further explored the clinical significance of TMB by comparing the percentage of patients with a high TMB (≥10 mut/Mb) and observed a higher percentage of TMB-high tumors in metastases from lung adenocarcinoma, HR+/HER2- ductal, lobular breast and prostate cancer patients. In five tumor types, we detected a significantly higher proportion of any actionable mutations (OncoKB levels 1 to 3, Methods) in metastases compared to primary tumors, but these differences were not significant after adjusting for differences in FGA and TMB (**Figure 2B; Table S2**). Next, we investigated differences in the frequency of arm-level copy number alterations between primary tumors and metastases. Because FGA was generally higher in metastases, we used a multivariable model to adjust for FGA and found 26 statistically significant differences (**Figure 2B; Table S2**). For example, in pancreatic adenocarcinoma, gain of chromosome 12p gain, where the oncogene *KRAS* is located, was more frequent in metastases than in primary tumors (17% vs. 4%, q-value = 0.002). In HR+/HER2- ductal and lobular breast cancer, loss of chromosome 16q, a feature of low-grade breast cancer (Natrajan et al., 2009), was more frequent in primary tumors than in metastases (41% vs. 30%, q-value = 1.56E-07 and 68% vs. 56%, q-value = 0.002, respectively). Finally, we investigated the frequency of recurrent oncogenic alterations between primary tumors and metastases and identified a total of 67 statistically significant differences across 17 tumor types. We also investigated the frequency of oncogenic pathways and identified 47 statistically significant differences across 11 tumor types (**Figure 2C; Table S2**). Amongst the statistically significant alterations, 53 were more frequent in metastases, while only 14 alterations were more frequent in primary tumors. *TP53* mutations were the most commonly observed significant alterations and were more frequent in metastases in 7 tumor types (lung adenocarcinoma, prostate adenocarcinoma, HR+/HER2- ductal breast, colorectal MSS, lobular breast cancer, pancreatic neuroendocrine and uterine endometrioid). A possible explanation is that *TP53* mutation is a later event in some of these tumor types; in others, it may simply be a hallmark of more aggressive disease. The notable exception was head and neck cancer, where *TP53* mutations were more frequent in primary tumors. Other genomic alterations that were most often enriched in metastases included *CDKN2A* deletion (significant in 5 tumor types), *PTEN* mutations and deletion (4 tumor types) and *MYC* amplification (4 tumor types). The most common significantly enriched oncogenic pathways in metastases were p53, Cell Cycle and DNA damage repair. The most significant differences were observed for alterations known to be associated with resistance to hormonal therapy in hormone-sensitive tumors. For example, *AR* amplification and *AR* mutations were significantly more frequent in prostate cancer metastases (1% vs. 30% and 0% vs. 6%, q-value < 0.05), and *ESR1* mutations were more frequent in HR+/HER2- ductal breast cancer (2% vs. 19%, q-value < 0.05), lobular breast cancer (2% vs. 13%, q-value < 0.05), and endometrioid uterine cancer metastases (3% vs. 10%, q-value < 0.05). These differences can likely be attributed to positive selection due to therapy since most patients with prostate cancer and ER+ breast cancer receive hormone therapy. *TERT* mutations were more frequent in metastases from papillary thyroid cancer and cutaneous melanoma patients (46% vs. 69% and 70% vs. 81%, q-value < 0.05), but higher in primary tumors from head and neck squamous cell carcinoma patients (41% vs. 25%, q-value < 0.05). *ALK* fusions, a predictive biomarker for the use of *ALK* inhibitors, were slightly more frequent in lung adenocarcinoma metastases (3% vs. 6%, q-value < 0.05). *KRAS* mutations were more frequent in metastases from pancreatic neuroendocrine patients (1% vs. 10%, q-value < 0.05) as was the overall frequency of RTK/RAS pathway alteration in this tumor type (6% vs. 21%, q-value < 0.05). While *KRAS* mutations are a hallmark of pancreatic adenocarcinoma, this could suggest the existence of a transdifferentiation mechanism from neuroendocrine to an adenocarcinoma phenotype during metastatic progression. Collectively, these data indicate that metastases have higher chromosomal instability across many tumor types and that mutations in a multitude of driver alterations occur at different frequencies in primary and metastatic tumors.

### Genomic differences between primary samples from metastatic and non-metastatic patients

Many of the primary tumors included in the previous analysis were from patients with metastatic disease. To identify genomic determinants of metastatic disease present in primary tumors we compared the genomic features of primary tumors from metastatic patients (n=11,993) to primary tumors from non-metastatic patients (n=3,669). The median follow-up time for these two groups was 33 months and 27 months, respectively. In 10 tumor types, FGA was significantly higher in primary tumors from metastatic patients as compared to primary tumors from patients without metastases. Compared to non-metastatic patients, TMB was significantly higher in six tumor types but lower in head and neck squamous cell carcinoma (**Figure S2A-B; Table S3**). When interrogating the frequencies of recurrent oncogenic alterations, we identified statistically significant frequency differences in 32 genes across 12 tumor types and 21 oncogenic pathways across 9 tumor types (**Figure S2C; Table S3**), with most observed at higher frequencies in primary tumors from metastatic patients. Compared to non-metastatic patients, *TP53* mutations were significantly more frequent in metastatic patients with lung adenocarcinoma, HR+/HER2- ductal breast cancer, lobular breast cancer, urothelial bladder cancer, prostate adenocarcinoma, and endometrioid uterine cancer. *TERT*promoter mutations were more frequent in metastatic patients with papillary thyroid cancer. The frequency of *MYC* amplification was significantly higher in metastatic patients with prostate adenocarcinoma, microsatellite stable (MSS) colorectal cancer, and TN ductal breast cancer. On the other hand, *SPOP* mutations were less frequent in primary tumors from metastatic prostate adenocarcinoma patients, *PIK3CA* mutations were less frequent in the primary tumors of HR+/HER2- ductal breast cancer metastatic patients, and *CDKN2A* mutations were less frequent in the primary tumors of pancreatic cancer metastatic patients. These findings support the hypothesis that a higher chromosomal instability is associated with metastatic progression in multiple tumor types and that several individual driver mutations might inform metastatic risk. Only a few of these, such as *SPOP* mutations in prostate cancer, which has been previously reported to be more frequent in primary tumors (Armenia et al., 2018), are associated with decreased metastatic potential.

### Genomic features associated with metastatic burden

To explore the genomic determinants of metastatic burden, we analyzed the relationship between genomic alterations and the number of metastatic sites per patient (n=21,546). Not surprisingly, a higher metastatic burden was significantly associated with shorter overall survival in most (39/50, 78%) tumor types (**Table S4**). We observed that chromosomal instability, as inferred by FGA, was positively correlated with metastatic burden on a pancancer level and in 11 individual tumor types. TMB, on the other hand, was not associated with metastatic burden on a pan-cancer level; it was positively correlated with metastatic burden in four tumor types, and negatively associated with metastatic burden in endometrioid and hypermutated uterine cancer (**Figure 3A-B; Table S5**). One of the strongest correlations between FGA and metastatic burden was observed in prostate cancer (rho = 0.33, q-value = 7.0E-45), which is in line with previous studies (Hieronymus et al., 2018; Taylor et al., 2010). Conversely, we did not observe such association in many tumor types, including MSS colorectal cancer, where chromosomal instability is already high in patients with low metastatic burden (**Figure 3B**).

**Figure 3.**
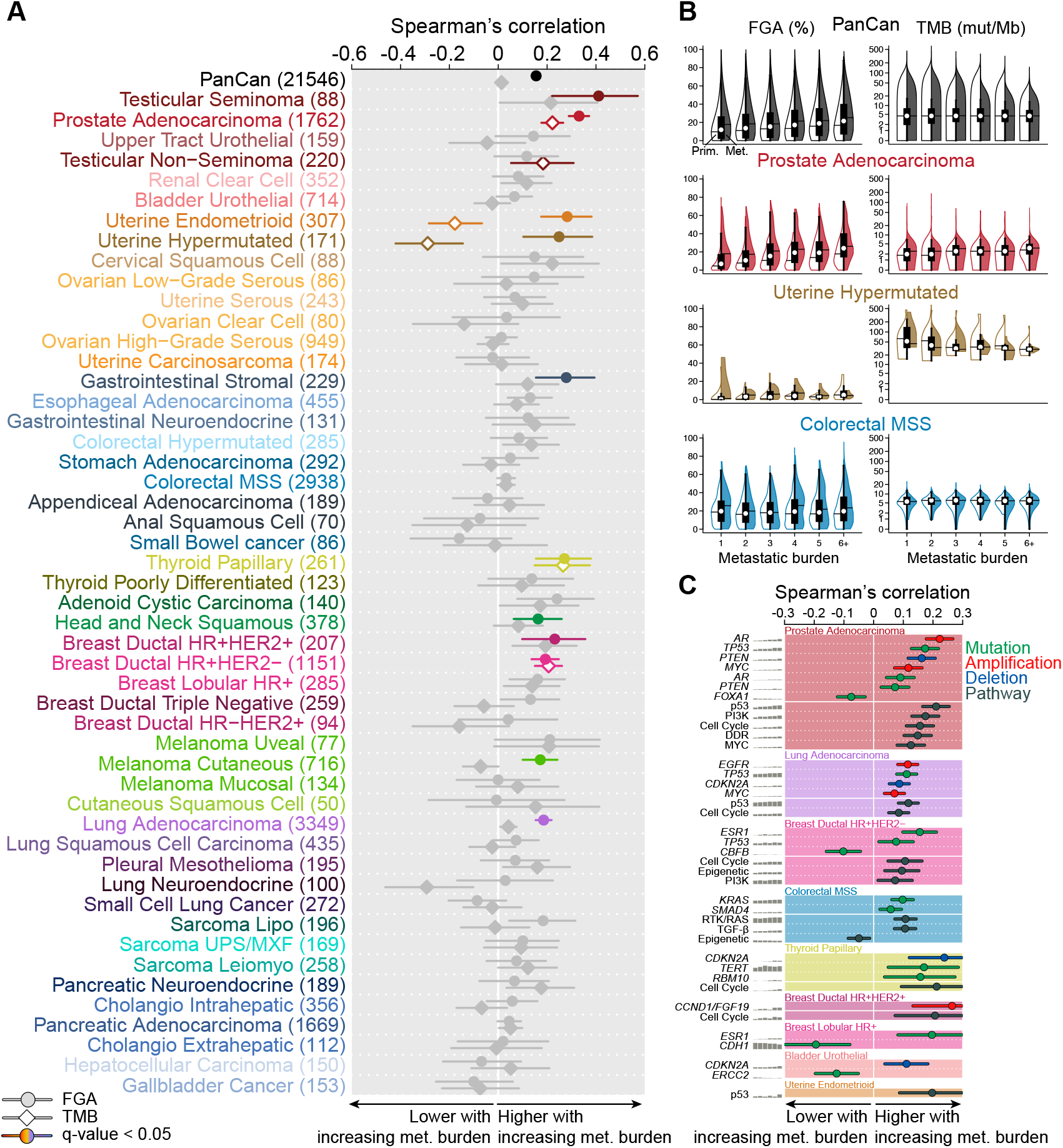
Genomic features associated with metastatic burden. (A) Spearman’s correlation coefficient between FGA (circle) and TMB (diamond) with metastatic burden. Associations without a significant trend are shown in grey, and the lines indicate 95% CI. (B) Correlation between FGA and TMB with metastatic burden in the entire data set, prostate adenocarcinoma, hypermutated uterine cancer, and MSS colorectal cancer. Boxplots display median point, IQR boxes and 1.5 × IQR whiskers for all samples. Split violin plots show the distribution of FGA and TMB in primary tumors (left, not filled) and metastases (right, filled). (C) Statistically significant oncogenic alterations and pathways associated with metastatic burden in individual tumor types. Spearman’s correlation coefficient is shown for each event, and the lines indicate 95% CI. Gene names in italics refer to specific genes, those in regular font refer to pathways. See also Table S4 and S5.

Next, we investigated the association between recurrent oncogenic alterations and metastatic burden and identified a total of 24 statistically significant associations across 8 tumor types. We also investigated the association with oncogenic pathways and identified 16 statistically significant differences across 7 tumor types (**Figure 3C; Table S5**). Consistent with its role as a gatekeeper against chromosomal instability (Bieging et al., 2014), we observed a significant positive correlation between *TP53* mutations and metastatic burden in prostate adenocarcinoma, lung adenocarcinoma and HR+/HER2- ductal breast cancer. There was also a significant positive correlation between p53 pathway alterations and metastatic burden in endometrioid uterine cancer. In metastatic prostate adenocarcinoma, *AR* amplification frequency was positively associated with metastatic burden. The frequency of *ESR1* mutations increased with metastatic burden in HR+/HER2- ductal and lobular breast cancer. *CDKN2A* deletion frequency was positively correlated with metastatic burden in bladder urothelial cancer, lung adenocarcinoma and papillary thyroid cancer, while *MYC* amplification frequency was associated with increasing metastatic burden in lung adenocarcinoma and prostate adenocarcinoma. Of note, the frequency of four oncogenic alterations and one oncogenic pathway were negatively correlated with metastatic burden; *FOXA1* in prostate adenocarcinoma, *CBFB* in HR+/HER2- ductal breast cancer, *CDH1* in lobular breast cancer, *ERCC2*in urothelial bladder cancer and the epigenetic pathway in MSS colorectal cancer (**Figure 3C; Table S5**). These results demonstrate that the relationship between higher chromosomal instability and increasing metastatic burden is tumor lineage dependent and that several driver mutations are associated with metastatic burden in both directions.

### Genomic differences of metastases according to their organ location

Next, we investigated the genomic characteristics of metastases (n=10,143) according to their organ location. As expected, the location of the sequenced metastases differed by tumor type (**Figure S3A**). We found 17 significant associations between FGA and the metastatic site in six tumor types, 10 of which were also significant when using adjusted FGA (**Figure S3B; Table S6**). CNS/Brain metastases from patients with lung adenocarcinoma, MSS colorectal cancer and cutaneous melanoma had a significantly higher FGA, while lymph node metastases from patients with lung adenocarcinoma, pancreatic adenocarcinoma, bladder urothelial and cutaneous melanoma had a significantly lower FGA. There were seven significant associations between TMB and the metastatic site. A total of 31 genomic alterations in nine tumor types were significantly associated with specific metastatic sites and 25 oncogenic pathways across six tumor types (**Figure S3C; Table S6**). *TP53* mutations were significantly more frequent in CNS/Brain metastasis from lung adenocarcinoma and liver metastasis from pancreatic adenocarcinoma, but less frequent in liver metastasis from urothelial bladder cancer and neuroendocrine lung cancer, as well as in intra-abdominal metastasis from pancreatic adenocarcinoma. In HR+/HER2- ductal breast cancer *ESR1* mutations were significantly more frequent in liver metastasis. In lobular breast cancer, *RHOA* mutations were significantly more frequent in ovarian metastasis and *FOXA1* mutations were enriched in liver metastasis. In lung adenocarcinoma, *CDKN2A* deletion was more frequent in skin and liver metastases but less frequent in lymph nodes. Similarly, in urothelial bladder cancer, *CDKN2A* deletion was more frequent in lung metastases but less frequent in lymph nodes. *PTEN* mutations, as well as PI3K pathway alterations, were higher in brain metastases from melanoma, which is in line with a previous melanoma-specific study (Bucheit et al., 2014). Among others, we found that *ERG* fusions were less frequent in bone metastasis of prostate cancer patients, *NF1* mutations were more frequent in lung metastasis of melanoma patients and that *FGFR3* mutations were more frequent in lung metastasis of bladder urothelial patients. Taken together, our results show that metastases from different organs can have different genomic makeup.

### Genomic features associated with metastasis to specific target organs

We analyzed the relationship between genomic features of metastatic patients and their organ-specific patterns of metastasis (n=21,546). We found 13 significant associations between FGA and organotropisms in 11 tumor types, seven of which were also significant when using adjusted FGA (**Table S7**). We observed a significant positive association between FGA and patients with liver metastasis in four tumor types (HR+/HER2- ductal breast, prostate adenocarcinoma, pancreatic adenocarcinoma and head and neck squamous), patients with lung metastasis in two tumor types (endometrioid uterine and cutaneous melanoma), and bonemetastasis in two tumor types (HR+/HER2- ductal breast and prostate adenocarcinoma). For TMB, we found eight significant associations between TMB and organ-specific patterns of metastatic in six tumor types, four positives (lung adenocarcinoma to brain and adrenal gland, pancreatic adenocarcinoma to liver, head and neck squamous to head and neck) and four negatives (prostate adenocarcinoma to bone, cutaneous melanoma to intraabdominal, lung adenocarcinoma to pleura and lung neuroendocrine to liver).

We found 57 significant recurrent oncogenic alterations associated with specific patterns of metastasis in 10 tumor types. When interrogating oncogenic pathway alterations, we found 48 significant associations in 12 tumor types (**Figure 4A; Table S7**). Lung adenocarcinoma, MSS colorectal cancer, and prostate cancer were associated with the highest number of significant associations. These results are summarized in **Figure 4B**. For example, lung adenocarcinoma patients with CNS/Brain metastasis had a higher frequency of *TP53* mutations, *TERT* amplification, and *EGFR* mutations, but a lower frequency of *RBM10* mutations. MSS colorectal cancer patients with lung metastasis had a higher frequency of *KRAS* mutations which was previously reported (Cejas et al., 2009; Pereira et al., 2015; Tie et al., 2011) but a lower frequency of *SRC* amplification. Prostate cancer patients with bone metastasis had a higher frequency of *AR* amplification and *PTEN* deletion but a lower frequency of *ERG* fusion; those with liver metastasis had a higher frequency of *PTEN* loss, *RB1* loss and *APC* mutations; those with brain metastasis had a higher frequency of *AR* amplification and NOTCH pathway alterations; and those with lung metastasis had a higher frequency of *APC* mutations and *CTNNB1* mutations. Experimental work has revealed the role of WNT pathway activation in driving prostate cancer metastasis (Leibold et al., 2020) and discovered a vulnerability to tankyrase inhibition in WNT altered prostate cancer. When interrogating the association between oncogenic pathways and organotropisms, we found that 26% of prostate cancer patients with lung metastasis had WNT pathway alterations, compared to 13% of patients without lung metastasis (**Figure4A**). As previously reported (Gerratana et al., 2020), *ESR1* mutations were more frequent in HR+/HER2- ductal breast cancer patients with liver metastasis (16% vs. 5%). *CBFB* mutations were less frequent in HR+/HER2- ductal breast cancer patients with bone metastasis, which was demonstrated in a mouse model (Ran et al., 2020) while alterations in the PI3K pathway were more frequent in patients with bone metastasis. HR+/HER2- ductal breast cancer patients with brain metastasis had a lower frequency of MAP3K1 mutations, which were recently shown to be a surrogate for the less aggressive luminal A breast cancer subtype (Nixon et al., 2019). In line with a previous study (Bucheit et al., 2014), *PTEN* mutations were more frequent in cutaneous melanoma patients with brain metastases, while *TP53* mutations were less frequent in those patients. Thyroid papillary cancer patients with bone metastasis had a higher frequency of *BRAF* mutations, and esophageal cancer patients with lung metastasis had a higher frequency of *ERBB2* amplification. In sum, while we did not observe gene or pathway alterations associated with specific routes of metastatic spread shared across different tumor types, our analysis revealed specific genomic alterations linked to specific organotropisms in individual tumor types.

**Figure 4.**
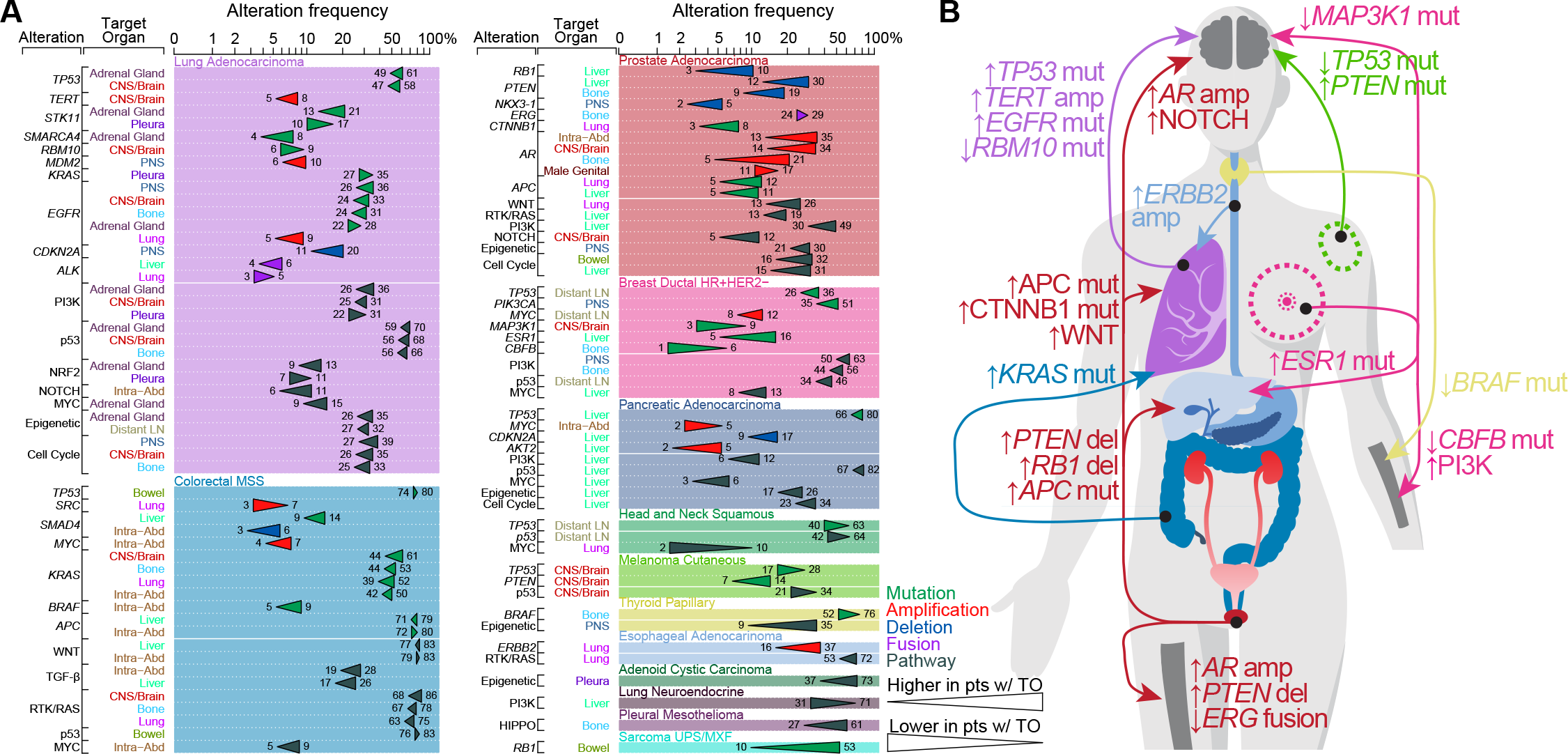
Genomic features associated with metastasis to specific target organs. (A) Statistically significant oncogenic alterations and pathways associated with organ-specific patterns of metastatic spread. Gene names in italics refer to specific genes, those in regular font refer to pathways. (B) Schematic drawing summarizing the main findings from (A). See also Table S7.

## DISCUSSION

We present MSK-MET, a unique, curated cohort of cancer patients with available genomic sequencing data and clinical information on metastatic disease and cancer outcome. Our study expands a previous pan-cancer dataset (Zehir et al., 2017) by including a larger number of patients with longer follow-up and by including a comprehensive description of metastatic events at the patient level. We demonstrate that mining of electronic health records can be used to extract relevant clinical information, and we present a pan-cancer map of metastasis in a contemporary cohort of patients treated at a single tertiary referral center.

Our analysis of genomic alterations from unpaired primary and metastatic samples revealed that metastases generally had a higher level of chromosomal instability, along with a higher frequency of WGD and *TP53* mutations. These results are consistent with previous studies that have shown an association between chromosomal instability and cancer progression (Bakhoum et al., 2018; Ben-David and Amon, 2020; Hieronymus et al., 2018; Shukla et al., 2020; Stopsack et al., 2019; Watkins et al., 2020). Our results also suggest that metastases generally have a higher fraction of clonal mutations. This lower intra-tumor heterogeneity could be attributed to clonal selection and selective pressure from cancer therapy (Birkbak and McGranahan, 2020). We also identified several genomic alterations and signaling pathways enriched in metastatic samples. As described before (Hu et al., 2020; Pareja et al., 2020; Razavi et al., 2018), the most significant enrichments were associated with known drug resistance mechanisms (e.g., *AR* alterations in prostate cancer, and *ESR1* mutations in breast cancer). We also compared primary tumor samples from metastatic and non-metastatic patients. In several tumor types, we observed a higher chromosomal instability and a higher frequency of *TP53* mutations amongst other drivers in primary samples from metastatic patients whereas the clonal fraction was generally similar. This highlights the importance of chromosomal instability in governing cancer progression and metastasis and shows that clonal selection is rather a hallmark of metastasis.

In an analysis aimed at identifying genomic alterations associated with metastatic burden, we found that higher chromosomal instability was correlated with metastatic burden in several tumor types. This association, however, was absent in many tumor types, including colorectal cancer, where copy-number alteration patterns may be established early in tumor development. Several mechanisms can explain the pro-metastatic effects of chromosomal instability and have been reviewed before (Ben-David and Amon, 2020). It is believed that chromosomal instability can promote tumor progression by increasing subclonal diversity and tumor evolution (Watkins et al., 2020), but aneuploidy itself is not a universal promoter of transformation and recent studies suggest that aneuploidy is cancer-type-specific (Ben-David and Amon, 2020), which is in line with our observations. Beyond global chromosomal instability, we also identified several specific genomic alterations and signaling pathways associated with metastatic burden. The majority, including alterations associated with drug resistance, were enriched in samples from patients with higher metastatic burden. Few were associated with lower metastatic burden, including *FOXA1* mutations in prostate cancer and *CBFB* mutations in breast cancer.

Lastly, we investigated associations between genomic alterations and specific routes of metastatic dissemination. We compared independent metastatic samples according to their organ sites. This revealed that the genomic landscape of metastasis differed according to their target organs. Previous studies have also interrogated the differences between primary tumors and metastatic sites using either independent samples (Armenia et al., 2018; Priestley et al., 2019; Robinson et al., 2017; Shih et al., 2020) or paired samples (Brastianos et al., 2015; Brown et al., 2017; Eckert et al., 2016; Hu et al., 2020; Jimenez-Sanchez et al., 2017; Makohon-Moore et al., 2017; Naxerova et al., 2017; Noorani et al., 2020; Reiter et al., 2020). Clinical data extraction from the EHR allowed us to explore the genomic alterations of metastatic patients by taking into consideration a greater part of the metastatic events occurring in a patient’s clinical course. We have generated a variety of hypotheses linking specific genomic alterations to specific organotropisms occurring in a cancer-specific manner. Future functional characterization of these alterations could result in the identification of novel biomarkers and therapeutic approaches for metastatic cancer.

Our study has several limitations. Firstly, while the overall cohort is large, sample size varied significantly between tumor types, which prevented us from drawing robust conclusions in less common tumor types. Secondly, the ICD billing codes used in our study likely do not fully capture all metastatic events and may be affected by inter-physician variability. Future improvements to the clinical data extraction process could come from the use of natural-language processing and machine learning approaches, which will be required to mine the wealth of data contained in EHR systems at scale. Despite these limitations, metastatic patterns observed across tumor types were consistent with previous reports (Budczies et al., 2015; Gao et al., 2019) and manual chart review of a subset of cases. Our study validated previous findings from others and was able to generate a vast array of new associations in several tumor types. The next challenge will be to prioritize the most clinically useful candidates to identify and validate prognostic and predictive biomarkers that will have the potential to influence the clinical management of patients.

Although this study represents a first step towards understanding how genomic alterations shape tumor progression, metastatic burden and organotropisms, more integrated studies are needed to fully investigate the impact of tumor cell-extrinsic effects, such as cancer therapy, target organ microenvironment and systemic factors. These studies will require comprehensive clinical timelines with accurate information about all lines of therapy and metastatic events. Additionally, single-cell profiling methods will be required to fully understand the cross-talk between tumor cells and the metastatic niche. Finally, our systematic study highlights the importance of chromosomal instability in progression and metastasis and drugs targeting this hallmark could represent an attractive strategy in several tumor types. MSK-MET will be publicly available via the cBioPortal for Cancer Genomics (Cerami et al., 2012; Gao et al., 2013) and will provide a valuable resource for the community and stimulate further research and applications in cancer care.

## ACKNOWLEDGMENTS

This study was supported by the MSK Cancer Center Support Grant (P30 CA008748), the MSK MIND initiative, Cycle for Survival, the Alan and Sandra Gerry Metastasis and Tumor Ecosystems Center, The Fund for Innovation in Cancer Informatics (ICI), the Robertson Foundation, a Prostate Cancer Foundation Challenge Award, and the Marie-Josée and Henry R. Kravis Center for Molecular Oncology.

## METHODS

### Samples and patients

A total of 43,400 solid tumor samples from 38,933 patients sequenced at Memorial Sloan Kettering Cancer Center from 2013-11-18 to 2020-01-06 (6.1y) and included in the AACR Project Genomics Evidence Neoplasia Information Exchange (GENIE) (AACR Project GENIE Consortium, 2017) 9.0-public database were considered for this study. All tumors were profiled using the Memorial Sloan Kettering Integrated Molecular Profiling of Actionable Cancer Targets (MSK-IMPACT) clinical sequencing assay, a hybridization capture-based, next-generation sequencing platform (Cheng et al., 2015). Tumor types were defined using a unique cancer type and one or more cancer type detailed (**Table S1**). For endometrial and colorectal cancers, we defined a subset of hypermutated (HM) tumors as those having an oncogenic *POLE* mutation or exhibiting more than 25 mutations/Mb or having MSIsensor score (Niu et al., 2014) > 10. Exclusion criteria were as follows: unavailable matched normal; low sequencing coverage (<100x); low tumor purity as defined by the absence of somatic alterations (including silent); pediatric patients (<18y at time of sequencing); patients with more than one unique sequenced tumor type; cancer of unknown primary; tumor type in which metastasis are rare (e.g. Gliomas); breast cancer with unavailable molecular subtype information; tumor types with small sample size (e.i. n <80 and either primary n <30 or metastasis n <30). Finally, one sample per patient was selected using a set of priority rules as follows: the presence of a FACETS fit that passed qc > highest purity > highest sample coverage > most recent gene panel. A total of 25,775 samples spanning 50 tumor types were used for analysis (**Figure S1A-D, Tables S1**). This set included samples that were sequenced with three generations of the MSK-IMPACT panel, containing 341 genes (n = 1,801 samples), 410 genes (n = 6,372 samples), and 468 genes (n = 17,602 samples).

### Clinical data extraction procedures for the identification and mapping of metastatic events

Clinical data were retrieved from the institutional electronic health records (EHR) database on 2020-11-05. Metastatic events were extracted from the pathology report of the sequenced samples and patients’ electronic health records. The anatomic location of the sequenced samples is described in the sample pathology reports as a free-text description by pathologists. The EHR includes International Classification of Diseases (ICD) billing codes which classify a comprehensive list of diseases, disorders, injuries and other health conditions including metastatic events. Metastatic events from the sample pathology report and the ICD billing codes from the EHR were systematically mapped to a curated list of 21 organs (**Table S8**). Lymph nodes were also classified as distant or regional given the anatomic location of the primary tumor. Of note, the classification of distant vs. regional was not possible for tumor types in which the anatomic location of the primary tumor is not well defined (e.i. melanoma cutaneous, cutaneous squamous cell, sarcoma lipo and sarcoma UPS/MXF). The organ site mapping for metastatic cancer is available at https://aithub.com/clinical-data-minina/oraan-site-mappina. For a user providing a table of organ site descriptions or ICD Billing codes, annotations of the 21 organ sites will be generated. Furthermore, additional annotations recognizing local extension and distant lymph node spread can be created. Metastatic burden was defined as the number of distinct organs (excluding regional lymph nodes) affected by metastases throughout a patient’s clinical course (ranging from 1 to 15 in the present study). Patients with more than six affected organ sites were grouped for analyses of metastatic burden.

### Comparison of metastatic sites automatically extracted from electronic health records vs. manual chart review

A total of 4,859 patients (22.5%) with metastatic sites extracted through manual chart review and previously published were available (Abida et al., 2017; Jones et al., 2021; Razavi et al., 2018; Shoushtari et al., 2021; Yaeger et al., 2018). Ten tumor types were represented including the most frequent (prostate, lung, breast, colorectal, and melanoma). There was a strong correlation between the number of metastatic sites retrieved from manual chart review and the number of metastatic sites automatically extracted from electronic health records (**Figure S1G**). For colorectal hyper mutant and MSS only the first metastatic events were reported so we restricted the comparison to the first metastatic event extracted from EHR. It is also important to note that the manual chart review was done before this study. Therefore, the present study has a longer follow-up which resulted in a higher number of metastatic sites. We also calculated the sensitivity for each metastatic site and each tumor type (**Figure S1H**). The median sensitivity was 77% across tumor types and metastatic sites.

### Genomic analysis

Tumor mutational burden (TMB) was calculated for each sample as the total number of nonsynonymous mutations, divided by the number of bases sequenced. Fraction of genome altered (FGA) was calculated for each sample as the percentage of the genome with absolute log2 copy ratios >0.2. Log2 copy-number ratios were derived as previously described (Cheng et al., 2015). Chromosome arm-level copy number alterations were computed using the ASCETS tool (Spurr et al., 2020) using default parameters. Allele-specific analyses of copy number alterations were performed using the FACETS tool (Shen and Seshan, 2016), which infers purity- and ploidy-corrected integer DNA copy number calls from sequencing data. The quality of FACETS fits was determined using a set of criteria as described in facets-preview (https://aithub.comAavlor-lab/facets-preview). To estimate a tumor purity- and ploidy-adjusted version of the FGA, we defined “adjusted FGA” as the fraction of the genome different from the major integer copy number (Mcn), where Mcn is defined as the integer total copy number spanning the largest portion of the genome. Tumor samples were considered to have undergone whole-genome doubling (WGD) if more than 50% of their autosomal genome had Mcn >2. The clonality of each mutation (clonal or subclonal or indeterminate) was determined as described in facets-suite (https://aithub.com/mskcc/facets-suite). For each tumor sample, the fraction of clonal mutations (clonal fraction) was determined by dividing the total number of clonal mutations by the sum of clonal and subclonal mutations. MSI-H status was defined by an MSIsensor score >10 (Niu et al., 2014). Somatic alterations were annotated using OncoKB for oncogenicity and clinical actionability (Chakravarty et al., 2017) (Data version: v2.8, released on 2020-09-17). For hypermutated colorectal and hypermutated uterine cancer, only genes that were recurrently mutated based on MutSig-CV (q-value<0.1) were considered for association analyses. For each tumor type, recurrent oncogenic alterations were defined as those considered oncogenic or likely oncogenic by OncoKB and present in at least 5% of either primary or metastatic samples (median of 15 per tumor type, **Table S1**). Canonical oncogenic pathway-level alterations were computed using curated pathway templates as previously reported (Ding et al., 2018; Sanchez-Vega et al., 2018). Segmented copy-number data were processed using the CNtools package v1.4.

### Statistical analyses

Comparisons between groups (primary vs. metastatic tumors, primary samples from metastatic vs. non-metastatic patients, and metastases according to their organ location) were performed using the non-parametric Mann-Whitney U test for continuous variables or the Fisher’s exact test for categorical variables. Differences in the frequency of actionable mutations (Levels 1 to 3, as defined by OncoKB) between groups (primary tumors vs. metastases and primary tumors from metastatic vs. non-metastatic patients) were further tested using a multivariable logistic regression model adjusted for TMB and FGA. Differences in the frequency of arm-level copy number alterations between groups (primary vs. metastatic tumors and primary samples from metastatic vs. non-metastatic patients) were tested using a multivariable logistic regression model adjusted for FGA. A genomic feature was considered to be significantly correlated with metastatic burden if (a) the Spearman’s correlation between the two variables was statistically significant (q-value < 0.05) and (b) the coefficient associated with the genomic feature as a predictive variable in a multivariable linear regression model adjusted for sample type (metastatic vs. primary tumor) was statistically significant (p-value < 0.05). The second condition was required because the ratio of metastatic samples to primary samples was associated with metastatic burden and could otherwise act as a confounding factor. We assessed genomic features associated with the presence or absence of metastasis in a target organ using only target organs present in at least 5% of the patients. A genomic feature was considered to be significantly associated with metastasis to specific target organs if (a) the Mann-Whitney U test for continuous variables or the Fisher’s exact test for categorical variables was statistically significant (q-value < 0.05) and (b) the coefficient associated with the genomic feature as a predictive variable in a multivariable logistic regression model adjusted for sample type (metastatic vs. primary tumor, categorical) and metastatic burden (1 to ≥6, numerical) was statistically significant (p-value < 0.05). The second condition was required because the ratio of metastatic samples to primary samples and metastatic burden were associated with metastasis to specific target organs and could otherwise act as a confounding factor. When TMB and FGA were used in a generalized linear model (linear and logistic model), their distributions were harmonized using a normal transformation as described before (Vokes et al., 2019) then scaled from 0 to 1 by subtracting the minimum and dividing by the maximum. Logistic regression was performed using Firth’s bias-reduction method as implemented in the R package brglm (Kosmidis and Firth, 2020). Overall survival (OS) was measured from the time of sequencing to death and was censored at the last time the patient was known to be alive. If a patient had more than one sequenced sample, the first time of sequencing was used.

Median follow-up time was calculated using the reverse Kaplan-Meier method. Median overall survival and five-year survival rate were calculated by the Kaplan-Meier method. The association between metastatic burden and overall survival was assessed using univariable Cox proportional hazards regression models. All reported p-values are two-tailed. Multiple testing correction was applied within each tumor type using the false discovery rate (q-value) method and q-value < 0.05 was considered significant. All analyses were performed using R v3.5.2 (https://www.R-project.org) and Bioconductor v3.4.

---

**Figure S1.**
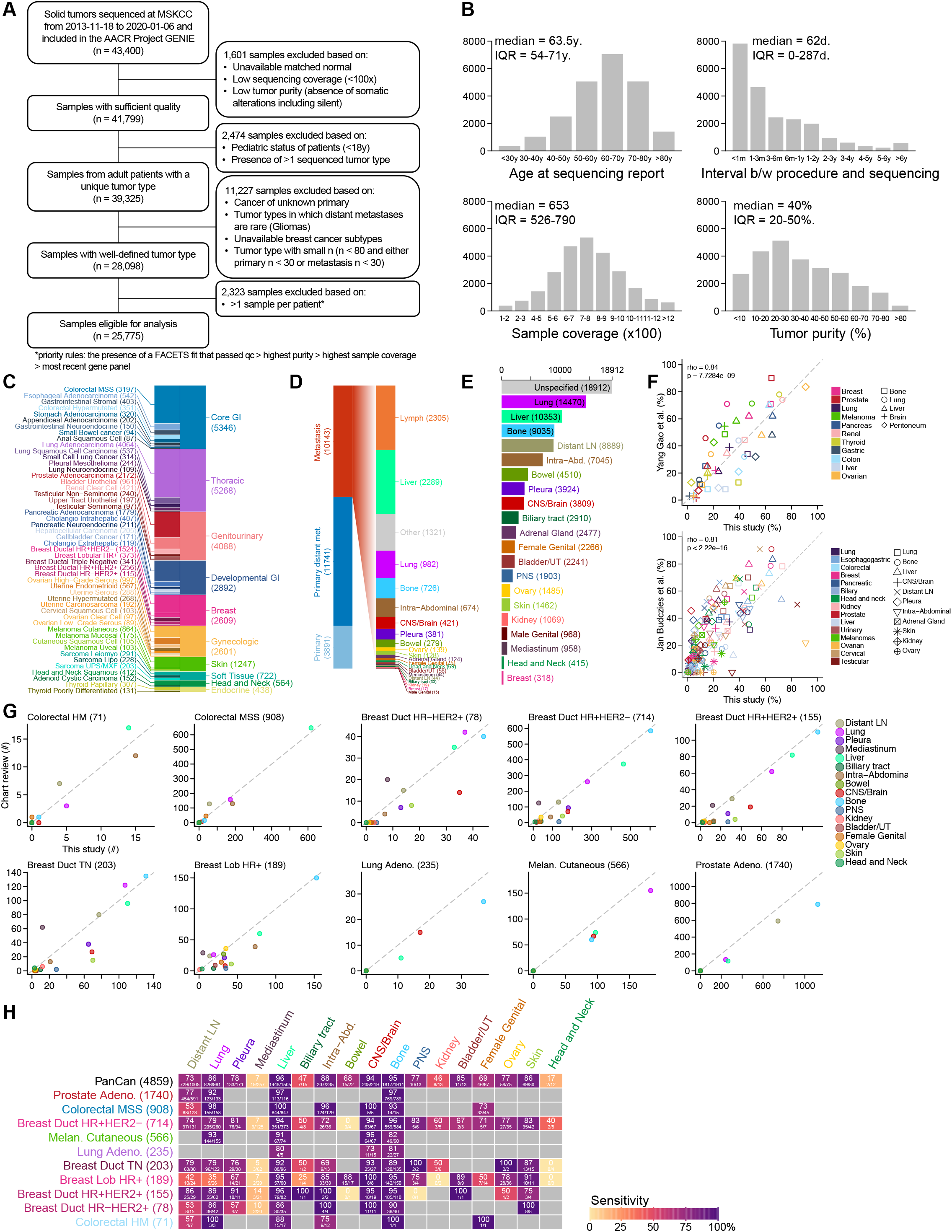
Study design and characteristics of the patients and samples included in MSK-MET, related to Figure 1. (A) CONSORT flow diagram of the study. (B) Distribution of age at time of sequencing, time interval between surgical procedure and sequencing, tumor sample coverage, tumor purity assessed by the pathologist. (C) Distribution of 25,775 tumors across 50 tumor types grouped by ten organ systems. (D) Distribution of the 25,775 tumors according to the sample type (primary vs. metastasis), site of metastatic sample and whether the primary sample was from a patient with evidence of distant metastasis at the time of the study or not. (E) Distribution of the 99,220 metastatic events mapped to 21 organ sites. (F) Comparison of the frequency of metastasis in several target organs from different tumor types reported in (Gao et al., 2019) and in (Budczies et al., 2015) vs. the present study. (G) Comparison of the number of metastasis using data from manual chart reviews and clinical data automatically extracted from the EHR (This study). (H) Heatmap showing the recall rate (sensitivity) across several target organs from different tumor types using patients retrieved from manual chart reviews.

**Figure S2.**
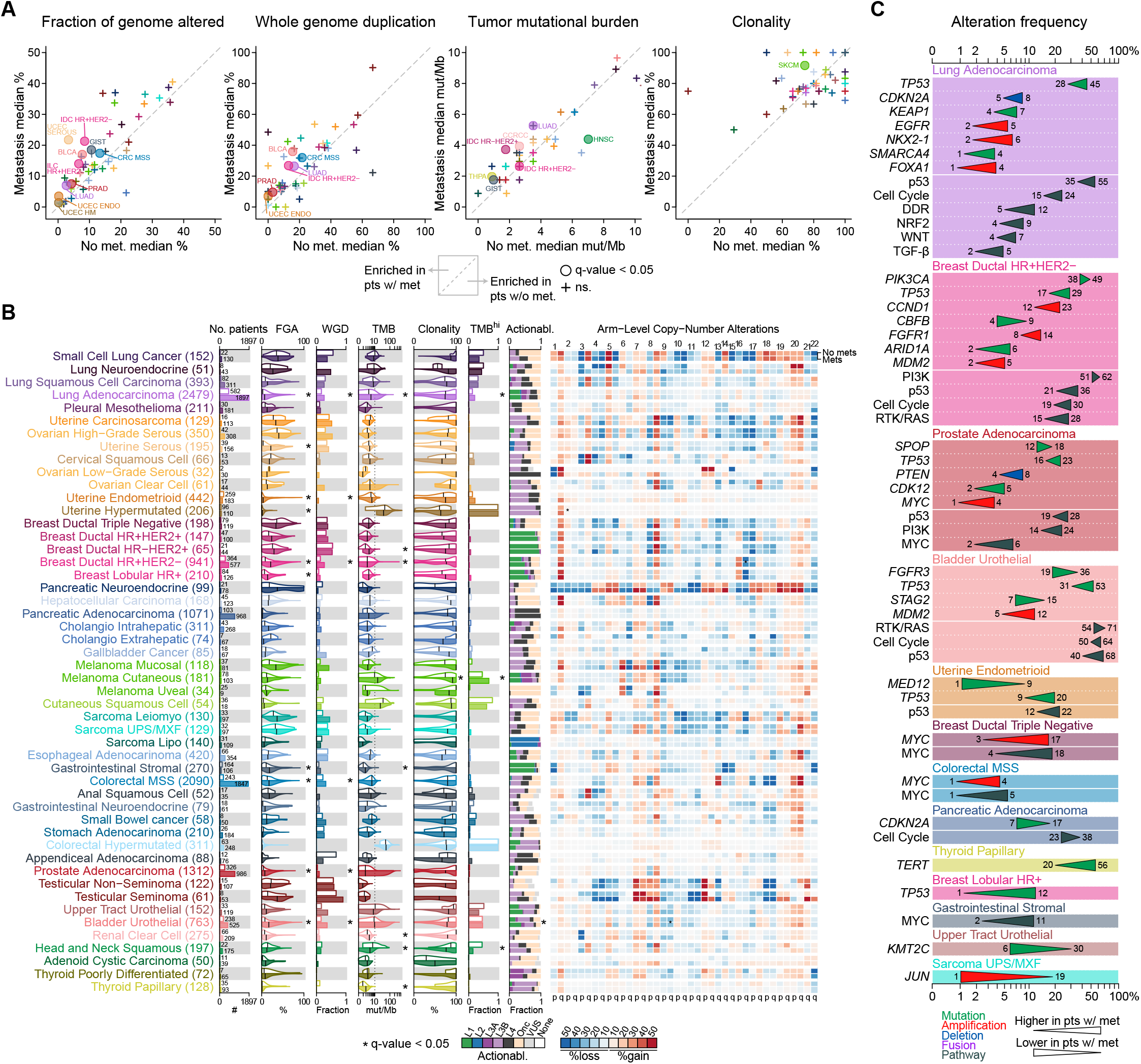
Genomic differences between primary samples from metastatic and non-metastatic patients, related to Figure 2. (A) Scatterplot showing the comparison of the median FGA, median WGD, median TMB and median clonality for each tumor type in primary samples from metastatic and non-metastatic patients. (B) The following clinical and genomic features are shown side-by-side for primary samples from metastatic and non-metastatic patients using a combination of barplots and violin plots; from left to right: sample counts, FGA, fraction of samples with WGD, TMB, clonality, fraction of samples with TMB-high status and distribution of highest actionable alteration. The black vertical line in each violin plots represents the median. Heatmap shows the frequency of arm level alterations in primary tumors and metastases. Tumor types are ordered from top to bottom by decreasing FGA in metastasis and grouped by organ systems. * indicates q-value < 0.05. WGD and clonality were available for a subset of 10,106 samples with FACETS data. (C) Statistically significant differences in the frequency of oncogenic alterations between primary tumors and metastases across all tumor types. Triangles summarize oncogenic alterations frequencies in primary samples from metastatic vs. non-metastatic patients and are colored according to alteration type.

**Figure S3.**
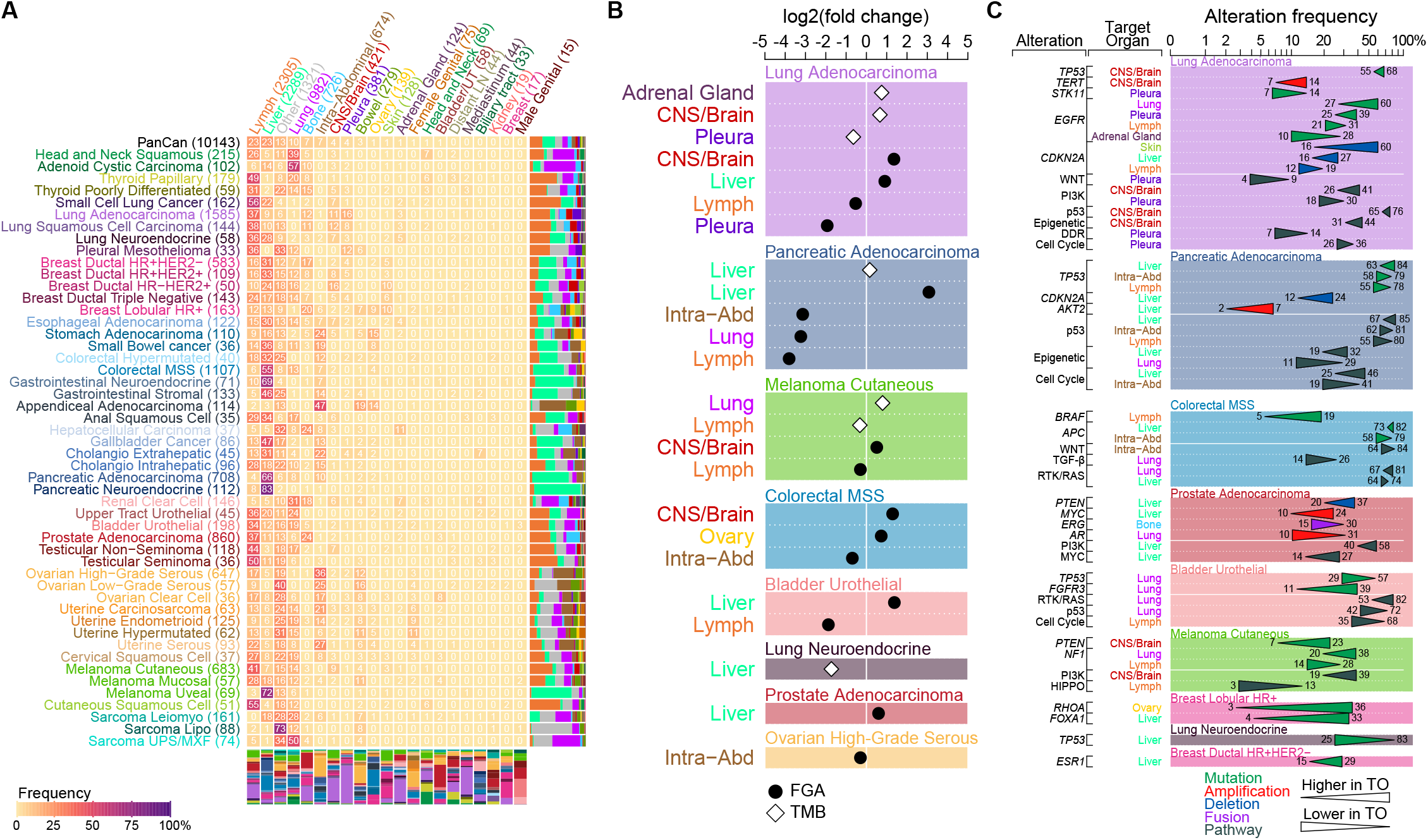
Genomic differences of metastases according to organ location, related to Figure 4. (A) Distribution of sequenced metastasis according to organ location with the heatmap showing the percentage of metastatic samples where each row represents a tumor type and each column represents the organ location. For each tumor type, the distribution of all 21 organ locations is shown as a stacked barplot to the right of the heatmap. For each organ, the distribution of all 50 tumor types is shown as a stacked barplot below the heatmap. For each tumor type, the number of metastasis samples is indicated in parentheses. For each organ, the number of metastasis samples is indicated in parentheses. (B) Statistically significant association between FGA (black circle) and TMB (white diamond) and specific metastatic sites. (C) Statistically significant oncogenic alterations associated with specific metastatic sites.

## SUPPLEMENTAL TABLES

Supplemental tables can be found online.

Table S1. Summary of the 50 tumor types included in the MSK-MET cohort, Related to Figure 1

Table S2. Genomic differences between primary tumors and metastases, Related to Figure 2

Table S3. Genomic differences between primary samples from metastatic and non-metastatic patients, Related to Figure S2

Table S4. Association between metastatic burden and overall survival assessed using Cox proportional hazards model, Related to Figure 3

Table S5. Genomic associations with metastatic burden, Related to Figure 3

Table S6. Genomic differences of metastases according to their organ location, Related to Figure S3

Table S7. Genomic features associated with metastasis to specific target organs, Related to Figure 4

Table S8. Mapping between free-text description from pathology reports and ICD billing codes from EHR to a curated list of 21 organs, Related to Figure 1

